# Inferring circadian phases and quantifying biological desynchrony across single-cell transcriptomes

**DOI:** 10.64898/2026.03.30.715278

**Authors:** Andrea Salati, Yves Paychère, Vincent Hahaut, Cédric Gobet, Felix Naef

## Abstract

Single-cell RNA sequencing (scRNA-seq) reveals heterogeneity in circadian clock states across individual cells, yet accurately inferring circadian phase and distinguishing biological desynchrony from technical noise remains challenging. Here, we introduce *scRitmo*, a probabilistic framework that infers single-cell circadian phases from mRNA count data, providing both a point estimate and a posterior uncertainty for each cell. A simulation-calibrated variance decomposition separates the observed phase dispersion into biological and technical components, enabling direct estimation of intercellular desynchrony. We validate *scRitmo* using deeply sequenced unsynchronized fibroblasts, where inferred transcriptomic phases accurately predict protein-level oscillations of a circadian reporter. Applied to murine scRNA-seq datasets from liver, aorta, and skin, *scRitmo* outperforms existing methods and reveals cell-type-specific levels of phase coherence. In SABER-FISH time-series data, the method recovers the progressive accumulation of desynchrony following synchronization, and in Drosophila clock neurons it captures cell-type-specific phase shifts and the expected increase in phase dispersion under constant darkness relative to light-dark entrainment. Together, *scRitmo* provides a principled approach for quantifying circadian (de)synchrony from transcriptomic data, decoupling biological phase variability from measurement noise across tissues, organisms, and experimental conditions.

## 2 Introduction

Circadian rhythms are evolutionarily conserved oscillations with a period of approximately 24 hours that coordinate physiology and behaviour with the solar day. In mammals, these rhythms are generated cell-autonomously and manifest in a transcriptional-translational feedback loop (TTFL) involving the core activators CLOCK and BMAL1, which drive the expression of their own repressors, *Periods* (*Pers*) and *Cryptochromes* (*Crys*) [41], with analogous mechanisms governing circadian rhythms in Drosophila [14].

The mammalian circadian system is organized hierarchically: the suprachiasmatic nucleus (SCN) acts as the master pacemaker and synchronizes peripheral tissue oscillators via neuronal and humoral cues [34]. Indeed, whole-brain imaging has revealed that circadian rhythmicity extends far beyond the SCN [50]. These peripheral oscillators are entrained by systemic signals such as feeding-fasting cycles [6] and daily body temperature rhythms [5]. Historically, gene expression rhythms in these peripheral tissues have been studied using bulk time-series tissue transcriptomics, which provides a population-averaged view of circadian gene expression parameters [52]. However, bulk approaches inherently obscure cell-to-cell variability in the underlying oscillatory gene circuits.

Pioneering studies using real-time bioluminescence reporters revealed that peripheral tissues and individual cells function as cell-autonomous, self-sustained oscillators that can exhibit various levels of phase desynchrony depending on the conditions, including initial synchronization and proliferation state [51, 45, 27, 46, 47]. This intercellular heterogeneity is a key feature of the system, arising from heritable genetic and epigenetic factors, as well as stochasticity in gene expression that set single-cell oscillatory properties such as the intrinsic periods [19, 20, 29]. This heterogenity is further shaped by cell-specific differences in sensitivity to entrainment cues, which ultimately dictate the individual phase of entrainment of each cell [30]. Furthermore, in organs such as the liver, single-cell gene expression heterogeneity also reflects a specific space-time logic where circadian phase is intertwined with spatial zonation [8]. More broadly, chronodisruption is itself an inherently multi-scale phenomenon, manifesting not only as misalignment between organ systems but also as phase incoherence among individual cells within a tissue [11].

Consequently, distinguishing true biological desynchrony, arising from heterogeneity in intrinsic period, sensitivity to entrainment, and spatial context, from technical noise is essential for understanding circadian organization in health and disease.

The advent of single-cell RNA sequencing (scRNA-seq) offers an unprecedented opportunity to dissect this heterogeneity at the transcriptomic level. To leverage this, several computational methods have been developed to infer biological phases or ordering from static single-cell snapshots. These range from oscillation detection algorithms such as Oscope [18] and reCAT [22], to autoencoder-based approaches such as Cyclum [21], spectral methods such as scPrisma [15], correlation-based tools such as tauFisher [10], and modelbased approaches such as TimeTeller [43]. Probabilistic frameworks such as Tempo [2] and VIST [49] have begun to address the stochastic nature of single-cell data. While these and other tools have advanced the field, accurately inferring circadian phase from scRNA-seq data remains challenging. A key distinction, relevant to all such methods, is between the *external time* at which a sample is collected (referred to as Zeitgeber time, ZT) and which typically refers to an entrainment signal, and circadian time (CT) reflecting the internal oscillatory state of each individual cell, which we refer to here as the (internal) *circadian phase*. It is this latter, cell-intrinsic phase, that *scRitmo* infers from gene expression data. The difference between external and internal phase is central in chronobiology as it depends on the entrainment and collective synchrony status, and underlies many key phenomena such as the chronotype in humans or internal desynchrony of organs or cells within a tissue. In this paper, we have chosen to report internal phases in units of time, i.e. using CT times between 0 and 24 h rather than less intuitive angles.

scRNA-seq data are inherently sparse and noise-prone. Due to limited capture efficiency, current protocols typically detect only a fraction (5–15%) of the endogenous transcriptome [40]. This is particularly consequential for core clock genes, many of which are transcription factors expressed at low copy numbers. Moreover, intrinsic biological fluctuations such as transcriptional bursting contribute substantially to expression variability, a phenomenon observed not only in cell culture but also in intact tissues [39, 13, 31].

Indeed, recent work using multiplexed SABER-FISH suggests that biological noise sets a fundamental limit to phase inference, with accurate single-cell phasing requiring signal aggregation from over 50 rhythmic genes, far more than the core clock circuit alone provides [28]. Without rigorous uncertainty metrics and a broader gene set, it is difficult to separate true biological desynchrony from technical noise sources [16].

To address these limitations, we introduce *scRitmo*, an unsupervised, probabilistic framework for single-cell circadian phase inference. *scRitmo* leverages a parametrized model of periodic gene expression, together with Negative Binomial noise modelling, to generate a posterior phase distribution for every cell, yielding both a point estimate and a measure of uncertainty. A key innovation is the ability to decouple biological phase variability from technical noise through a simulation-calibrated variance decomposition that recovers the true biological desynchrony (*σ*_bio_) from the observed data. We validate *scRitmo* across diverse biological contexts, data modalities, and organisms, demonstrating that it enables the quantitative study of circadian (de)synchrony across scales, from individual cells to tissue-level organization.

## 3 Results

### 3.1 *scRitmo*: A probabilistic framework for circadian phase and desynchrony estimation

To address the aforementioned challenges in circadian scRNA-seq analysis, we developed *scRitmo*, an unsupervised probabilistic framework for inferring single-cell circadian phases and quantifying biological desynchrony (Fig. 1A) in cell populations. Unlike methods that provide only point estimates, *scRitmo* models single-cell count data using a Negative Binomial (NB) distribution and computes a full posterior phase distribution for each cell, *P* (*θ*|**x**), where *θ* represents the circadian phase and **x** the observed gene expression vector. This yields both a phase estimate (taken here as the posterior mode) and a measure of uncertainty, defined as the circular standard deviation (cSTD) of the posterior *σ*_*u*_. Periodic gene expression profiles are modelled as harmonic oscillations, defining the expected mean expression levels used to compute the NB likelihoods for the gene counts. The model parameters (gene harmonic coefficients, dispersion coefficients) are learned by maximizing the marginal likelihood after integrating out the latent cell phases (Methods). A reference clock model (peak phases and amplitudes) for a set of *Clock-Reference* genes was derived from harmonic regressions using bulk time-series RNA-seq data from multiple organs (Fig. 1B, Fig. S1A) [52]. These *Clock-Reference* genes were selected based on established roles in the circadian literature and significant rhythmicity across the majority of profiled tissues (Methods). While the model implementation allows for the optimization of all gene parameters including peak phases (*φ*_*g*_), we found that fixing these parameters based on established biological knowledge often provides a useful constraint that stabilizes inference, preserving the interpretability and reference of the inferred cell phases. Notably, this constraint preserves the known phase relationships of genes such as *Bmal1* and *Per2*, ensuring a robust mapping of the circadian phase despite the sparsity of scRNA-seq data.

**Figure 1:**
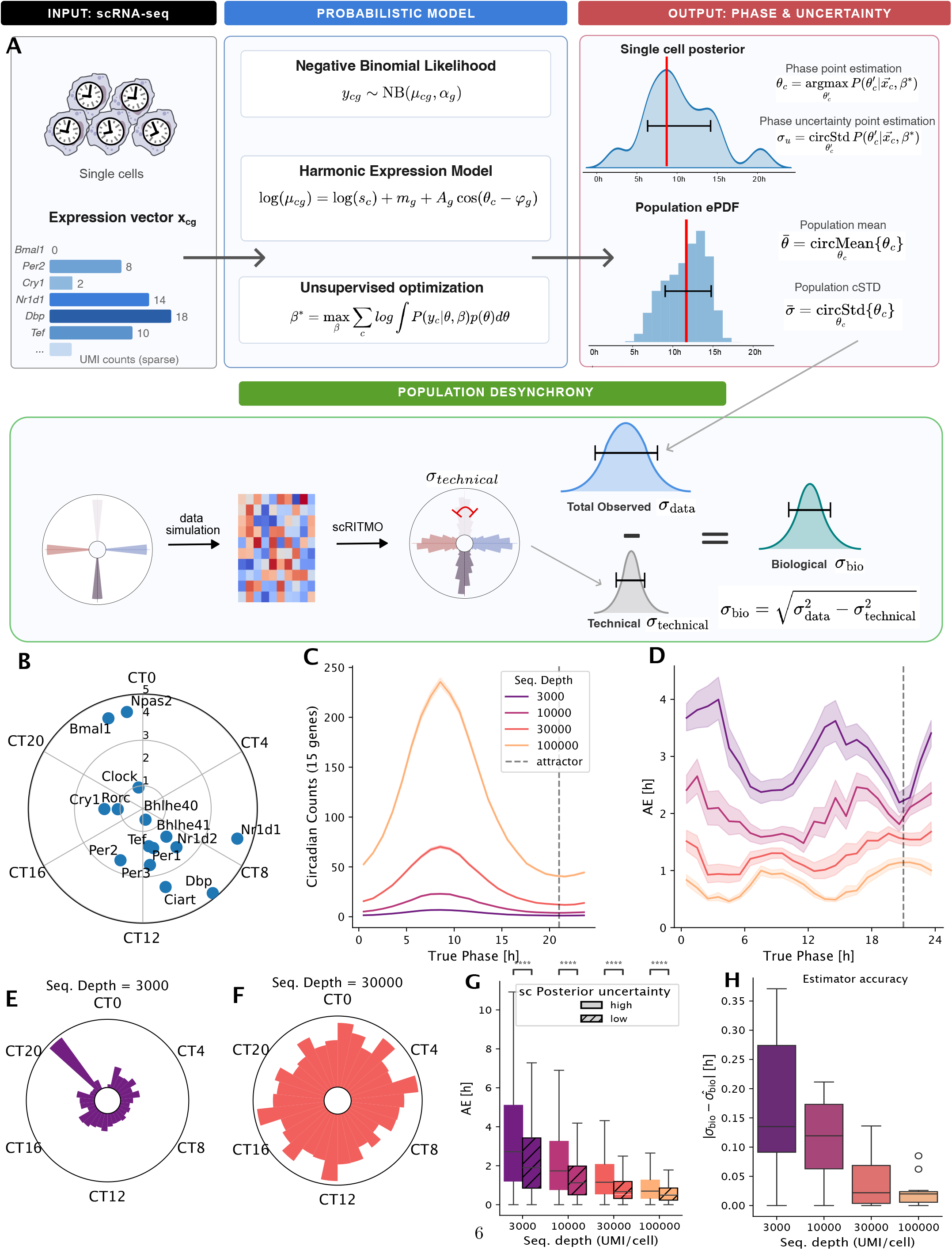
*scRitmo*: A probabilistic framework for circadian phase and desynchrony estimation. **A**. Schematic overview of the *scRitmo* framework. The pipeline uses a raw count matrix of circadian genes, such as *Bmal1* and *Per2*, as input. The probabilistic model describes gene counts using a negative binomial likelihood parameterized by a harmonic expression function and normalizing by the library size *s*_*c*_. Through unsupervised optimization, the method infers a posterior distribution for each cell, yielding a phase point estimate (*θ*_*c*_) and a phase uncertainty (*σ*_*c*_), which are aggregated to form the population phase distribution. The lower section illustrates the quantification of population desynchrony, where the total observed circular standard deviation (*σ*_*data*_) is decomposed to separate technical noise (*σ*_*technical*_) from the underlying biological variance (*σ*_*bio*_). **B**. Polar plot of peak phases and amplitudes (log2 fold change) for *Clock-Reference*, estimated by fitting a consensus harmonic model across organs from bulk time-series RNA-seq data [52]. **C–H. Simulation-based benchmarks. C**. Total “circadian counts” (summed expression of 15 genes in *Clock-Reference*) as a function of true phase (simulated) across sequencing depths. Dashed line indicates the attractor phase. **D**. Median Absolute Error (MAE) of phase inference on simulated data across the 24h cycle at varying sequencing depths. **E– F**. Polar histograms of inferred phases from a ground-truth uniform distribution at low (**E**, 3’000 UMI/cell), and high (**F**, 30’000 UMI/cell) sequencing depth. **G**. Phase inference error stratified by posterior uncertainty. Simulated cells are stratified into below-median (hatched) and above-median (solid) uncertainty groups based on the posterior cSTD. ****: *P <* 0.0001 (Mann-Whitney U test), *n* = 5000 simulated cells. **H**. Accuracy of the *σ*_*bio*_ estimator across sequencing depths. Each point represents an independent simulation with a unique ground-truth *σ*_*bio*_. The y-axis shows the resulting absolute error 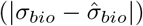.

To benchmark *scRitmo* under controlled conditions, we designed simulations to generate synthetic scRNA-seq datasets with known ground truth circadian phases using *Clock-Reference* gene profiles. We define the total circadian counts as the summed transcript counts from the set of *Clock-Reference* genes, which strongly varies along the 24h cycle (Fig. 1C). A major peak occurs around ZT10, where several genes, *Dbp, Tef, Nr1d1* (Rev-Erb*α*), *Nr1d2* (Rev-Erb*β*), and *Ciart* reach their expression maxima. Conversely, the *Bmal1* phase region corresponds to a trough where circadian counts are approximately 4-fold lower. At low sequencing depths (e.g., 3,000 UMI/cell), this non-uniform expression density creates a “phase attractor” where inferred phases artificially cluster, a bias that is progressively mitigated as depth increases (Fig. 1D–F, Fig. S1B–D). These results suggest that achieving robust estimates requires either high sequencing depth (*>* 20, 000 UMI/cell) or incorporating a broader set of phase-telling genes.

A central feature of *scRitmo* is its estimation of phase uncertainty. We examined the correspondence between the posterior phase uncertainty *σ*_*u*_ and the actual inference error, quantified as the absolute error between simulated ground truth phases and inferred phases (Methods). In simulated datasets, cells with low posterior uncertainty exhibited a reduced absolute error compared to those with high uncertainty; Fig. 1G), indicating that the model’s internal confidence is linked with accurate inference at the single-cell level. We next addressed the challenge of separating biological from technical phase variability in populations of cells. The stochasticity inherent in Negative Binomial count sampling artificially inflates the circular variance of inferred phases: even a perfectly synchronized population will exhibit apparent phase dispersion due to measurement noise alone. To decouple these sources, we developed a variance decomposition strategy (Fig. 1A). For a given dataset, *scRitmo* first estimates the observed dispersion of inferred phases across cells (*σ*_data_). It then quantifies the expected (background) technical contribution (*σ*_technical_) by performing inference on simulated data that matches the dataset’s sequencing characteristics but assumes a perfectly synchronized population. The true (residual) biological desynchrony is then calculated as: 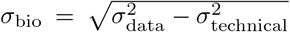 (Fig. 1A, Fig. S1E). We observed that *σ*_technical_ is not uniform around the clock but varies significantly across true phase (Fig. S1F), reflecting the non-uniform distribution of peak phases across the rhythmic gene set. This leads to increased uncertainty in regions where fewer genes are peaking or counts are low, such as near the *Bmal1* trough (Fig. S1G).

Benchmarking this estimator across simulated sequencing depths shows that the absolute error of 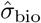 is remarkably low (under 0.2h) even at the lowest sequencing depths (Fig. 1H). By decoupling biological from technical variance, *scRitmo* provides a principled approach for quantifying cell-to-cell desynchrony directly from transcriptomic data.

### 3.2 *scRitmo*-inferred phases align with protein levels and capture progressive desynchrony in 3T3 fibroblasts

To validate that *scRitmo*-inferred phases reflect true circadian time, we profiled 695 unsynchronized 3T3 fibroblasts using plate-based scRNA-seq (MERCURIUS™ FLASH-seq), generating a multimodal dataset with simultaneous FACS-based quantification of a RevErb*α*-Venus fluorescent circadian reporter [27] (Fig. 2A–E). The plate-based protocol provides high sequencing depth and sensitivity per cell (850k median reads/cell).

**Figure 2:**
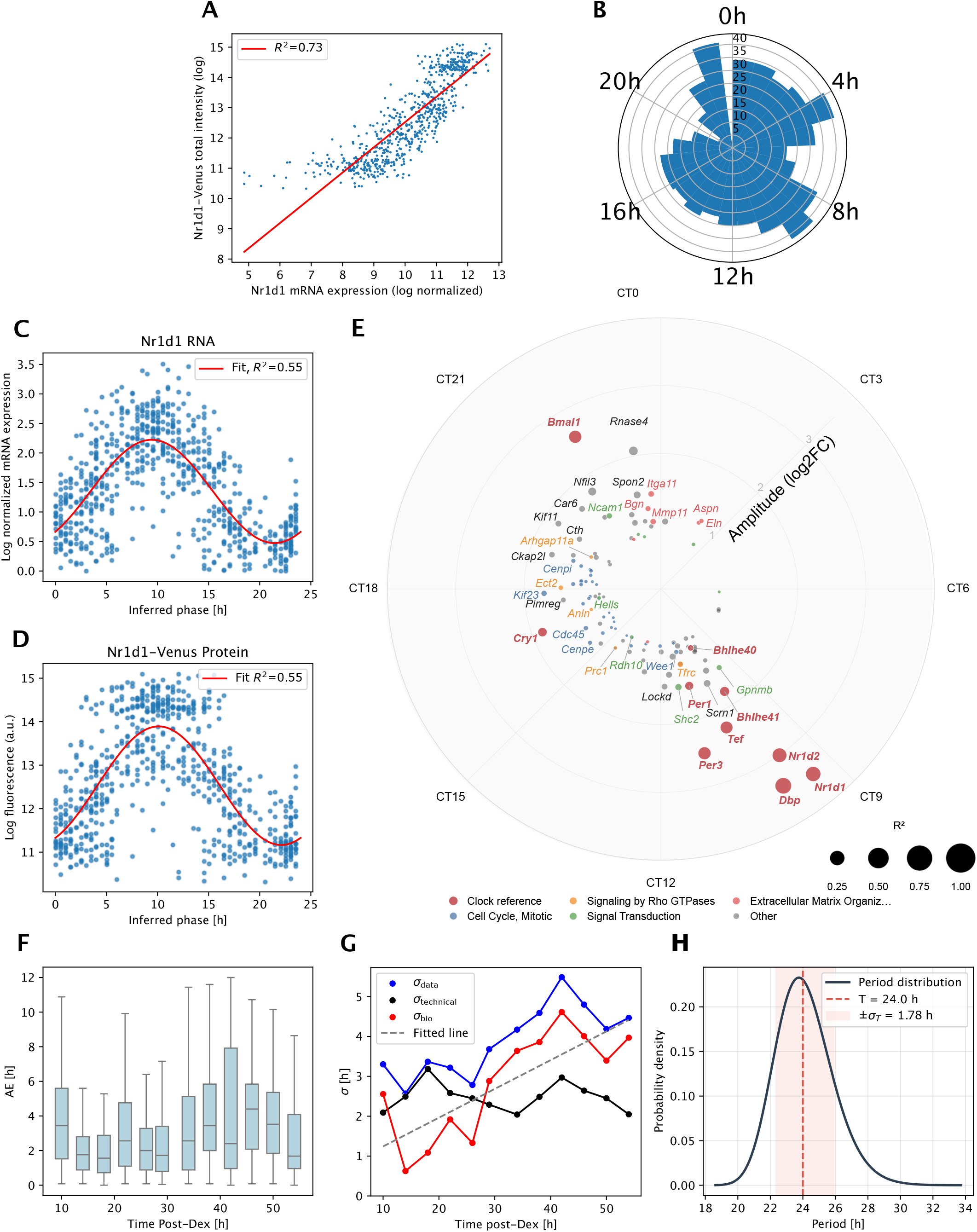
*scRitmo*-inferred phases align with protein levels and capture progressive desynchrony in 3T3 fibroblasts. **A–E**. Analysis of plate-based scRNA-seq (FLASH-seq) data in unsynchronized 3T3 fibroblasts with simultaneous FACS-based quantification of a Nr1d1-Venus (Rev-Erb*α*-YFP) circadian reporter. **A**. Scatter plot showing the relationship between normalized and log-transformed endogenous *Nr1d1* mRNA against total fluorescence intensity at 530nm wavelength (variance explained: *R*^2^ = 0.73). **B**. Circular histogram of inferred cellular phases in the unsynchronized culture. **C–D**. Normalized *Nr1d1* mRNA counts (C) and Rev-Erb*α*-YFP protein fluorescence (D) plotted against the phase inferred from the scRNA-seq data. The red line represents a harmonic fit to the data (*R*^2^ = 0.55). **E**. Polar plot displaying genes identified as significantly rhythmic via genome-wide harmonic regression against *scRitmo*-inferred phases. The angular coordinate represents the peak phase of expression in circadian time (CT), while the radial distance indicates the oscillation amplitude (log_2_ fold change). The size of each point is proportional to the coefficient of determination (*R*^2^) of the harmonic fit. Colors indicate functional categories and Reactome pathway membership: clock reference genes (red), cell cycle and mitotic processes (blue), signaling by Rho GTPases (orange), signal transduction (green), and extracellular matrix organization (pink). Representative genes such as *Wee1, Prc1, Mmp11*, and *Nfil3* are labeled. **F–H**. Analysis of progressive desynchronization in SABER-FISH time-series data post-dexamethasone (Dex) shock in 3T3 fibroblasts [28]. **F**. Boxplots of the MAE of the phase inference over time (10–55h post-Dex). **G**. Biological desynchrony (*σ*_bio_, red) and technical noise (*σ*_technical_, black) over time (post-dex). **H**. Estimation of the underlying distribution of single-cell free-running periods based on the rate of desynchronization (slope of *σ*_bio_ in G). The solid black line is the mapped period distribution, red dotted the 24h period line and the shaded area corresponds to *±σ*_*T*_ = 1.78*h*

We first confirmed the correspondence between the transcriptome and the reporter signal, observing a high correlation between endogenous Rev-Erb*α* (*Nr1d1*) mRNA levels and (Rev-Erb*α*)-YFP protein fluorescence intensity (*R*^2^ = 0.73, Fig. 2A). Given the high sequencing depth of FLASH-seq and simulation results at high coverage, phase inference was performed using the *Clock-Reference* gene set alone (Fig. S2A). The inferred phases were broadly distributed across the 24h cycle, consistent with the unsynchronized state of the cells (Fig. 2B). The inferred phase accurately reconstructed the oscillation of both the endogenous (Rev-Erb*α*) mRNA level and the (Rev-Erb*α*)-YFP protein fluorescence (Fig. 2C, D), showing a clear rhythmic alignment between the *YFP* transcript and its protein across the inferred cycle (Fig. S2B). The good fit for the reconstructed protein cycle (*R*^2^ = 0.55) provides evidence that *scRitmo* infers a phase that is not only transcriptionally accurate but relevant at the reporter protein level.

To assess whether the inferred phases capture biologically meaningful temporal structure beyond the input genes, we performed a genome-wide harmonic regression against the *scRitmo*-inferred phases. This analysis identified over 100 significantly rhythmic genes beyond the *Clock-Reference* (Fig. 2E). Pathway enrichment analysis (Reactome [26]) revealed that these genes were enriched in cell cycle regulation (*Wee1, Cenpe*), Rho GTPase signaling (*Prc1, Tfrc*), extracellular matrix organization (*Mmp11, Aspn*), and broader signal transduction pathways (*Shc2, Ncam1*). The recovery of established circadian output genes absent from the reference set, such as *Nfil3, Wee1, Tfrc*, together with the coherent enrichment of clock-regulated processes, confirms that *scRitmo* infers phases reflecting genuine circadian phase. We next tested whether *scRitmo* could track the temporal evolution of phase dispersion using SABER-FISH time-series data from dexamethasone (Dex)-synchronized 3T3 fibroblasts [28] (Fig. 2F–H). In this longitudinal design, cells were sampled at intervals from 10 to 55 hours post-synchronization as they naturally drifted out of phase, providing a controlled setting in which desynchrony is expected to increase monotonically over time. This dataset represents a different challenge for phase inference, as it contains a limited set of four clock reference genes (*Bmal1, Nr1d1, Nr1d2, Tef*). However, the high sensitivity provided by SABER-FISH (Fig. S2C) allowed *scRitmo* to perform with remarkable precision, yielding biologically correct phase relationships for the four genes (Fig. S2D). Despite the minimal gene set, the model achieved a median absolute error (from ZT time) of approximately 2.2 hours during the initial 10–34h period and 3.1h during the later time points (33–55h) (Fig. 2F).

*scRitmo* also captured an increase in the posterior uncertainty *σ*_*u*_ over the course of the experiment, suggesting that the molecularly defined circadian phase becomes less certain with time after the initial synchronization stimulus (Fig. S2E).

More importantly, *scRitmo* captured the expected progressive increase in phase dispersion. By applying the variance decomposition, we separated measurement error from true biological heterogeneity and observed that while *σ*_technical_ remained approximately constant over time, *σ*_bio_ increased linearly from 10 to 55 hours post-Dex (Fig. 2G). Notably, the earliest time point (10h) showed elevated MAE and *σ*_bio_ (Fig. 2F,G), consistent with incomplete synchronization immediately following the Dex pulse. The slope of this increase indicates that the circular standard deviation grows by approximately 0.1 hours (5.4 minutes) per elapsed hour, directly reflecting the rate at which individual cellular oscillators drift apart due to differences in their intrinsic periods and other noise sources.

This linear relationship is consistent with theory and allows estimating the underlying distribution of single-cell free-running periods. Indeed, assuming cell-specific frequencies that are fixed over time and modelling each cell’s phase evolution deterministically as *φ*(*t*) = *ωt*, where angular velocities *ω* are normally distributed, the population’s circular standard deviation grows as cSTD(*t*) = *σ*_*ω*_ · *t* [27]. The observed slope thus provides a direct estimate of *σ*_*ω*_, from which we derived the probability density of single-cell periods via a reciprocal transformation (Fig. 2H). The variance of the resulting period distribution (*σ*_*T*_ = 1.78*h*) closely matches values reported from longitudinal time-lapse fluorescence and luciferase imaging studies [46, 29, 27], demonstrating that *scRitmo* can recover the intrinsic dispersion of cellular clock speeds from static population snapshots. This measurement is traditionally accessible only through continuous single-cell tracking.

### 3.3 Robust phase inference with expanded phase-telling gene sets reveals cell-type specific desynchrony in mice

To evaluate *scRitmo* in complex biological contexts, we applied our framework to murine scRNA-seq time-series datasets from liver [8], aorta [2], and skin [9], encompassing diverse cell types including hepatocytes, smooth muscle cells (SMC), fibroblasts, and tissue-resident immune populations. These datasets exhibited varying levels of sequencing depths across the profiled populations, with liver hepatocytes exhibiting the lowest depth in our study (~2,000 UMI/cell) (Fig. S3A).

As expected from our simulation predictions, phase inference based solely on the *Clock-Reference* in hepatocytes exhibited a pronounced bias, with cell phases artificially clustering around the region of low counts (Fig. 3A). To overcome this, we followed two paths. First, we identified cell-type-specific *Extended-Set* s using a pseudobulk approach, selecting genes by rhythmic significance and amplitude from harmonic regressions against the known collection time (see Methods). This strategy expands the predictor set from the 15 *Clock-Reference* to approximately 70–100 high-confidence rhythmic genes, comprising a mixture of clock- and system-driven transcripts (Supplementary Table 1 contains the rhythmic genes parameters for each cell type). In hepatocytes, the *Extended-Set* successfully eliminated the attractor bias and significantly reduced the Median Absolute Error (MAE) (Fig. 3B). We further evaluated an alternative gene set meant to be the most faithful with respect to the core-clock machinery, comprising genes identified as rhythmic in wild-type mice but arrhythmic in *Bmal1* and *Cry1/2* double-knockout models [44] (Fig. 3C). This knockout-validated set comprising 70 genes achieved comparable accuracy to the *Extended-Set*, demonstrating that when curated biological knowledge is available, *scRitmo* can achieve high precision without relying on external time labels for gene selection.

**Figure 3:**
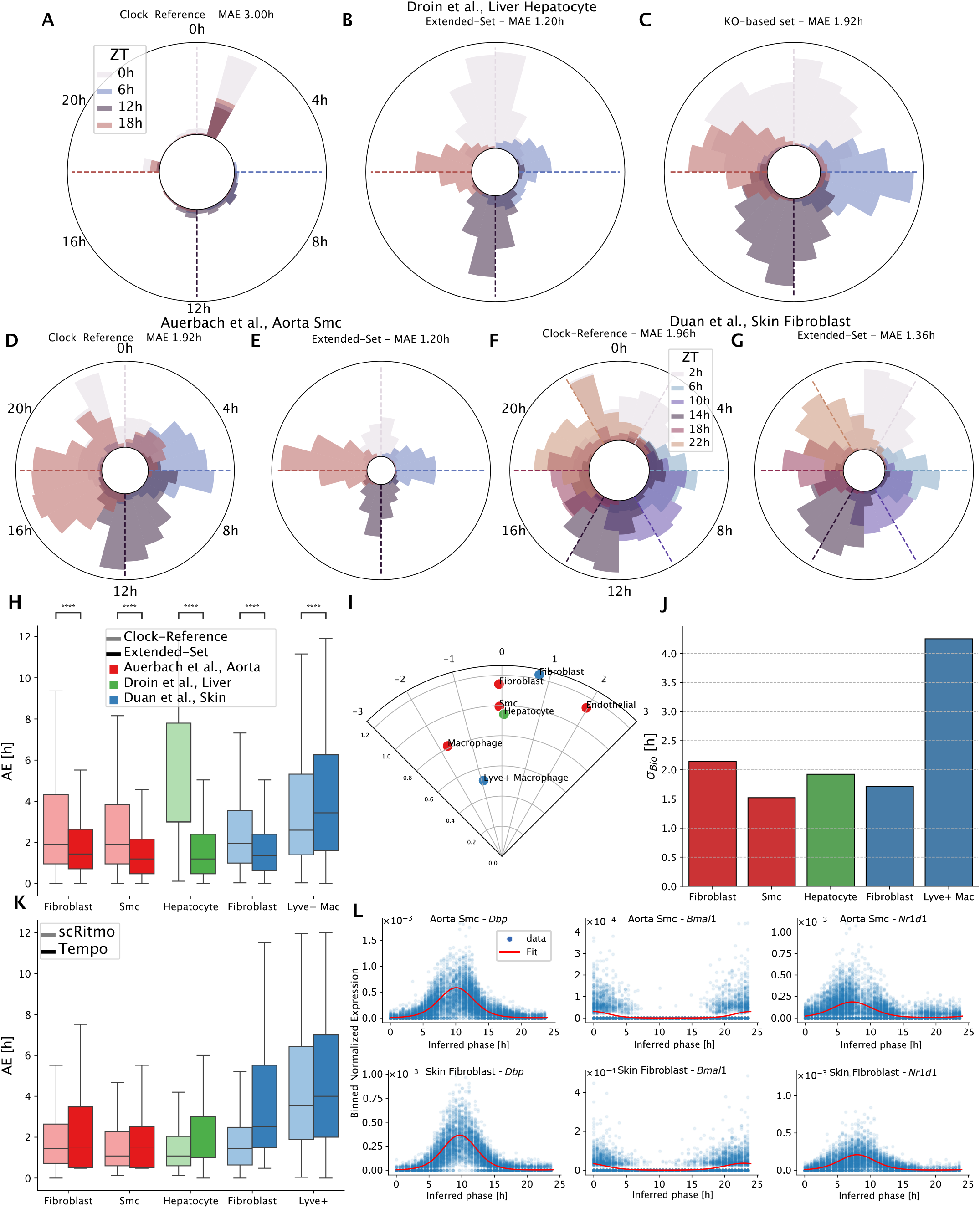
Robust phase inference with expanded phase-telling gene sets reveals cell-type specific desynchrony in mice. **A–G**. Circular histograms (rose plots) of inferred cellular phases across different tissues and reference gene sets. Different colors represent distinct external collection times; all samples were collected under standard light-dark (LD, ZT) conditions. **A–C**. Liver hepatocytes ([8]) analysed using the *Clock-Reference* (A) already displayed in (Fig. 1B), an *Extended-Set* including food-driven rhythmic genes (B), and genes rhythmically expressed in WT but arrhythmic in *Bmal1* and *Cry1/2* double knockout mice [44] (C). **D–E**. Aorta Smooth Muscle Cells (SMC) [2] analysed with the *Clock-Reference* (**D**) and the *Extended-Set* (**E**). **F–G**. Skin Fibroblasts [9] analysed with the *Clock-Reference* (**F**) and the *Extended-Set* (**G**). **H**. Boxplots of the estimated MAE [h] for each cell type, comparing accuracy between *Clock-Reference* (light color) and the *Extended-Set* (dark). **I**. Polar visualization of global clock properties. The angle represents the phase delay between cell types, while the radius corresponds to the average amplitude of *Clock-Reference*. **J**. Quantification of biological desynchrony. Bar plots show the estimated circular standard deviation (*σ*_*Bio*_) indicative of genuine biological heterogeneity within each cell type. **K**. Benchmarking against Tempo. Boxplots comparing the MAE distributions of *scRitmo* versus Tempo using the *Extended-Set*. **L**. Gene expression dynamics. Normalized expression of representative core clock genes (*Dbp, Bmal1, Nr1d1*) plotted against *scRitmo*-inferred phases in Aorta SMC (top) and Skin Fibroblasts (bottom), with fitted harmonic regression lines (red).

Similar improvements were observed in aorta SMC and skin fibroblasts (Fig. 3D–G). In both populations, the cell-type-specific *Extended-Set* provided a less biased phase distribution and significantly lower MAE compared to the *Clock-Reference* alone (Fig. 3H). Notably, *σ*_*u*_ was substantially higher for *Clock-Reference* compared to *Extended-Set* (Fig. S3B).

Together, these results confirm that aggregating rhythmic signals from a broader set of transcripts provides the robustness necessary for precise single-cell phasing at the sequencing depths typical of droplet-based scRNA-seq.

Stratifying cells by posterior uncertainty *σ*_*u*_ effectively separated accurate from high-error phase inferences across all tissues (Fig. S3C, E, F). Notably, *σ*_*u*_ outperformed library size as a predictor of inference error (Fig. S3D), confirming that posterior uncertainty captures the quality of the circadian signal in each cell rather than reflecting global sequencing depth. Beyond phase accuracy, *scRitmo* enables the quantification of biological desynchrony at the tissue level. Polar visualizations revealed distinct phase relationships between cell types, with radii corresponding to the average amplitude of *Clock-Reference* genes (Fig. 3I). Notably, *Lyve1* ^+^ skin macrophages displayed a much weaker *Clock-Reference* expression.

Across all cell types, we estimated *σ*_bio_ using the *Extended-Set* for phase determination (Fig. 3J). For fibroblasts, smooth muscle cells, and hepatocytes, *σ*_bio_ ranged between approximately 1.1-1.6 hours, indicating that peripheral clocks maintain relatively tight phase coherence under standard light-dark entrainment. Such residual biological phase heterogeneity most likely reflects the equilibrium between intracellular noise driving phase diffusion and systemic entrainment cues acting as an external synchronizing force, resulting in a coherent but spread phase distribution.

In head-to-head benchmarking, *scRitmo* outperformed the probabilistic framework Tempo [2] across cell types on the *Extended-Set*, achieving lower MAE distributions while maintaining significantly lower runtime (Fig. 3K, Fig. S3G).

Reordering cells by scRitmo-inferred phase recovers coherent oscillatory profiles for core clock genes (*Dbp, Bmal1*, and *Nr1d1*) in both aorta SMC and skin fibroblasts (Fig. 3L). Red curves show harmonic regression fits to the phase-ordered expression, and the expected gene-specific peak times are recovered in both cell types, confirming that the inferred phases reflect genuine circadian structure.

*Lyve1* ^+^ macrophages exhibited the highest estimated desynchrony (*σ*_bio_ ≈ 4.2 h); however, unlike the other cell types, the *Extended-Set* for this population yielded higher MAE than the *Clock-Reference* alone (Fig. 3H), suggesting that non-circadian transcripts distort the inferred phases and the estimated dispersion.

Finally, we explored the limits of circadian organization in immune cells within the aorta and skin. Skin *Lyve1* ^+^ macrophages and aortic macrophages, despite exhibiting weaker oscillatory amplitudes than non-immune cell types (Fig. 3I), displayed canonical rhythmic signatures: their *Clock-Reference* genes maintained the expected phase relationships and reached statistical significance for rhythmicity. In contrast, other immune populations such as dendritic cells and T/NK cells exhibited disrupted or arrhythmic patterns (Fig. S4A–E), with substantially higher posterior uncertainties *σ*_*u*_ consistent with incoherent core clock expression (Fig. S4F).

### 3.4 Circadian desynchrony increases under free-running conditions in *Drosophila* clock neurons

To investigate how light-entrainment shapes the precision of circadian timekeeping at singlecell resolution, we applied *scRitmo* to scRNA-seq data of *Drosophila* clock neurons profiled under Light-Dark (LD) and Constant Darkness (DD) conditions across six time points [24]. This dataset, generated using an optimized CEL-Seq2 protocol, comprises ~1850 deeply sequenced cells spanning 16 clock neuron clusters (Fig. S5A,B), including the main s-LNv, l-LNv, LNd, DN1p, DN2 populations and subtypes [33]. This system provides a well-characterized setting in which light acts as a synchronizing Zeitgeber, and its removal is expected to increase inter-individual and inter-cellular phase dispersion.

We first characterized the *Drosophila-Set*, identified across cell types, for phase inference (Methods). This gene set includes canonical *Drosophila* clock and clock-controlled genes (*tim, per, vri, Clk, cry, Pdp1, cwo*), known output genes (*Argk1, CNMa*), and several previously uncharacterized rhythmic transcripts (*CG15628, CG32369*, among others) (Fig. 4A, Supplementary table 2). Together, these genes span the full circadian cycle with distributed peak phases and varying amplitudes (Fig. 4A), and cover a broad dynamic range in mean expression (Fig. S5C). The aggregate circadian transcript counts display clear temporal oscillations across most cell types (Fig. S5D).

**Figure 4:**
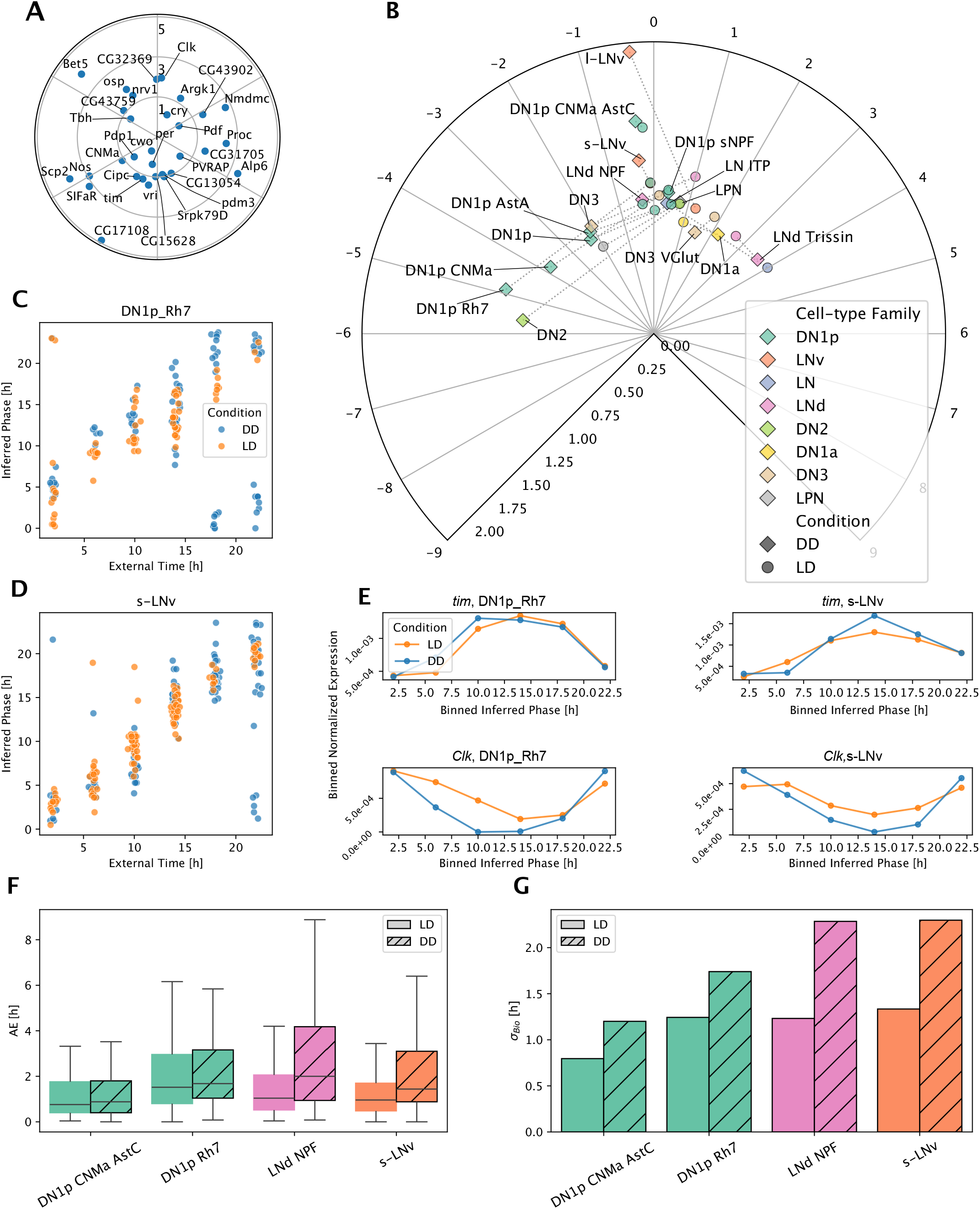
Circadian desynchrony increases under free-running conditions in *Drosophila* clock neurons. **A**. Polar plot displaying the global properties of the selected pan-rhythmic genes (*Drosophila-Set*), with angle representing the circular phase and radius representing the amplitude. **B**. Polar plot of average cellular phases per clock neuron cluster. Angular position encodes the mean phase offset; radial distance encodes the average amplitude of the cluster (Methods). Colours indicate cell-type family; circle and diamond markers distinguish Light-Dark (LD) and Constant Darkness (DD) conditions, respectively. **C, D**. Inferred circadian phase versus external time for the DN1p Rh7 (**C**) and s-LNv (**D**) clusters under LD (orange) and DD (blue) conditions. Jitter is added along the x-axis for visibility. **E**. Expression profiles of core clock genes *tim* (top) and *Clk* (bottom) in DN1p Rh7 (left) and s-LNv (right) clusters under LD (orange) and DD (blue) conditions. Inferred phases are binned into six intervals, and normalized expression values are averaged within each bin **F**. Boxplots of the phase absolute error (AE) for major clusters (*n >* 130 cells) under LD (solid) and DD (hatched) conditions. **G**. Estimated biological desynchrony (*σ*_bio_) for major clusters under LD (solid) and DD (hatched) conditions.

Projecting the mean inferred phase for each clock neuron cluster revealed systematic phase shifts between conditions (Fig. 4B). Under DD, most clusters exhibited phase advances relative to their LD counterparts. The magnitude of the shift varied markedly across neuron types, with DN1p Rh7 displaying the largest phase advance while the s-LNv and l-LNv pacemaker neurons showed minimal or no shift, consistent with their established role as the principal drivers of free-running rhythms (Fig. 4C, D). Since the dataset was collected at DD3 (i.e., three days into constant darkness), the large phase advances observed in some DN1p clusters suggest that acute network reorganization upon light removal, rather than slow period-driven drift alone, contributes to the observed phase redistribution. These phase shifts are reflected at the gene expression level: in DN1p Rh7, the expression profiles of *tim* and *Clk* are shifted between LD and DD, whereas in s-LNv they remain aligned across conditions (Fig. 4E). Oscillation amplitudes remained comparable after phase reassignment across most clusters (Fig. 4B), indicating that the cell-autonomous oscillatory machinery is preserved and that the primary effect of light removal is a reorganization of phase rather than a dampening of rhythmic output (Fig. 4E).

We next assessed inference accuracy using the Absolute Error (AE) across major cell types with respect to the external time (Fig. 4F). Phase inference was consistently more accurate under LD conditions, with DD datasets yielding higher estimation errors. The single-cell phase uncertainty *σ*_*u*_ also increased in DD conditions (Fig. S5E).

To determine whether this increased uncertainty reflects genuine biological heterogeneity rather than a technical limitation, we applied the variance decomposition to estimate *σ*_bio_ (Fig. 4G). Across the selected major clusters, biological desynchrony was markedly higher under DD than under LD entrainment. Importantly, because each time point consists of pooled neurons dissected from multiple animals, the observed *σ*_bio_ primarily captures interindividual variation in clock phase rather than within-brain cell-to-cell desynchrony. In DD, the absence of a shared light cue removes the common synchronizing signal, allowing inter-individual phase differences, and potentially inter-cellular ones, to accumulate through differences in intrinsic period and network-level reorganization. Interestingly, the magnitude of the LD-to-DD increase in *σ*_bio_ varied across cell types (Fig. 4G), suggesting that different clock neuron populations differ in their sensitivity to photic entrainment, in the strength of their coupling to the light-responsive pacemaker network, or in the tightness of intercellular coupling within each cluster.

## 4 Methods

### 4.1 *scRitmo* core model

*scRitmo* models the observed single-cell count matrix *X* ∈ ℕ ^*C×G*^, where *x*_*cg*_ represents the read count for gene *g* in cell *c*. Each cell is governed by a latent circadian phase *θ*_*c*_ ∈ [0, 2*π*). To account for the sparsity and overdispersion inherent in scRNA-seq data, counts are modelled using a Negative Binomial (NB) distribution:

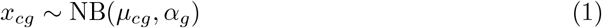

where *µ*_*cg*_ is the expected mean expression level and *α*_*g*_ is a gene-specific dispersion parameter. The log-mean expression is defined as:

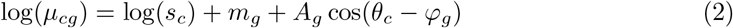

Here, *s*_*c*_ is the library size (total counts per cell), *m*_*g*_ is the basal expression level (mesor) of gene *g, A*_*g*_ is the amplitude, and *φ*_*g*_ is the acrophase (peak phase). We restrict the model to a single harmonic, as including higher-order terms increased complexity without improving fit and led to overfitting at the typical single-cell sequencing depths.

#### Parameter estimation via marginal likelihood maximization

*scRitmo* builds on Tempo’s strategy [2] of treating cell phase as a nuisance parameter and marginalizing it out. The two frameworks differ in two key respects. First, Tempo uses stochastic variational inference to approximate a posterior distribution over gene parameters, whereas *scRitmo* directly maximizes the marginal likelihood via gradient descent, yielding point estimates for the gene parameters *β*. Second, Tempo’s objective involves a marginalization of the loglikelihood weighted by a grid-based posterior *P* (*θ* | *β*, **x**_*c*_) that must be updated iteratively, whereas *scRitmo* integrates over the phase domain under a fixed uniform prior 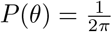. The gene-specific parameters 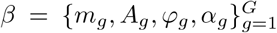 are estimated by maximizing the marginal likelihood obtained by integrating out the latent cell phases. Jointly optimizing both *β* and Θ is computationally unstable. Assuming conditional independence of genes given the phase, the marginal likelihood for cell *c* is:

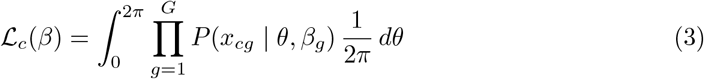

This integral is approximated numerically: we discretize the phase domain [0, 2*π*) into *M* equidistant points {*ϑ*_1_, …, *ϑ*_*M*_} and approximate the integral using Simpson’s rule or a summation approximation:

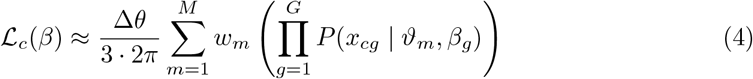

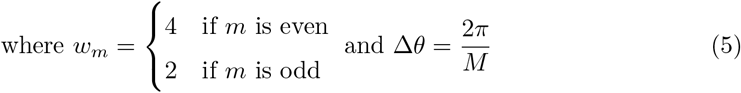

with *M* = 24 during training and *M* = 100 at evaluation.

The model parameters *β* are estimated by maximizing the log-marginal likelihood over all cells:

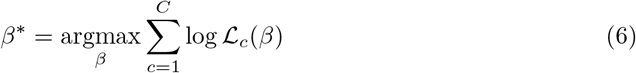

Optimization is performed using stochastic gradient descent (Adam optimizer) with default hyperparameters (*β*_1_ = 0.9, *β*_2_ = 0.999) in PyTorch. Gene-specific mesor parameters were initialized to their empirical values, while amplitudes were initialized to zero to prevent spurious rhythmic patterns during early optimization. To ensure biological interpretability of the inferred cellular phases, gene-specific peak times (*φ*_*g*_) were treated as fixed parameters across most experiments (the determination of *φ*_*g*_ is addressed in the the next section).

#### Training schedule

To ensure convergence across cell types with varying number of cells, the number of training epochs was adapted to dataset size: 1,200 epochs with batch size 32 for small datasets (*n <* 500 cells), 600 epochs with batch size 128 for medium datasets (500 ≤ *n <* 3,000), and 150 epochs with batch size 128 for large datasets (*n* ≥ 3,000). Training can be performed on GPU for accelerated computation.

#### Phase inference and uncertainty quantification

Once the gene parameters *β*^∗^ are learned, the circadian phase of each cell is inferred from the posterior distribution:

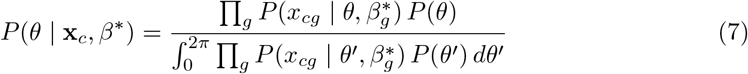

This posterior distribution provides a complete description of the phase estimate, capturing the uncertainty implied by the NB noise model Fig. 1A. *scRitmo* defines the phase point estimate *θ*_*c*_ as the maximum a posteriori (MAP):

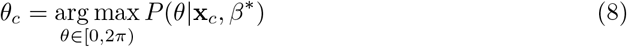

The associated uncertainty is quantified by the circular standard deviation (cSTD) of the posterior:

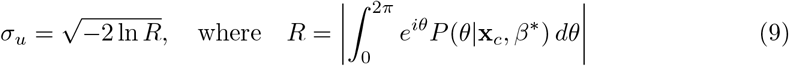

is the mean resultant length. Both the point estimate *θ*_*c*_ and the uncertainty *σ*_*u*_ are linearly scaled from radians to circadian hours using the factor 24*/*2*π*.

### 4.2 Simulations and desynchrony estimation

To simulate biologically accurate circadian expression profiles, the Fourier parameters from *Clock-Reference* were used. Cells were sampled from a Negative Binomial whose mean expression is modeled by Eq. (2).

#### Simulation of controlled desynchrony

To calibrate the relationship between observed and true phase dispersion, we generated synthetic scRNA-seq data with known biological desynchrony. Simulated cell phases were drawn from a Von Mises distribution:

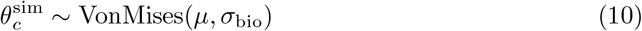

where *σ*_bio_ is the ground-truth biological desynchrony (parameterized via circular standard deviation). Synthetic count vectors were then sampled from the trained generative model:

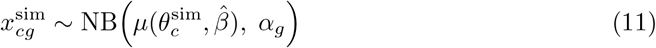

using the expression model defined in Eq. (2). Re-applying *scRitmo* to these synthetic transcriptomes yielded inferred phases whose circular standard deviation, *σ*_data_, consistently exceeded *σ*_bio_ due to technical noise introduced by the Negative Binomial sampling process (Fig. S1E).

#### Variance decomposition

We observed that the total observed phase variance decomposes additively (Fig. S1E):

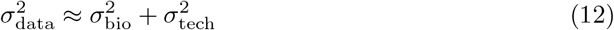

where *σ*_tech_ is the technical desynchrony, defined as the output phase spread obtained from a simulation with perfectly synchronized input phases (*σ*_bio_ = 0). This decomposition yields a corrected estimator for the biological component:

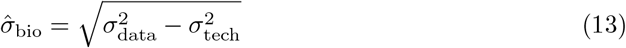

Both *σ*_data_ and *σ*_tech_ vary with circadian phase (Fig. S1F,G), introducing a phase-dependent bias. To obtain robust estimates across conditions composed of multiple batches, we aggregated variances using weighted means:

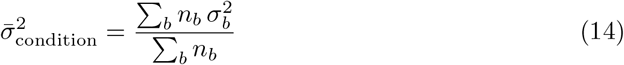

where *n*_*b*_ is the number of cells in batch *b*. The final 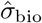 was computed by subtracting the aggregated technical variance from the aggregated total variance. In real data *σ*_bio_ is unknown. This procedure is applied for each biological replicate, and to estimate properly *σ*_tech_ the library sizes of real cells were used to generate the synthetic data.

### 4.3 Construction of reference gene sets

#### Consensus reference clock (*Clock-Reference*)

To establish a reference clock across multiple organs, we used a context-dependent extension of the core model. The log-mean expression for context *k* (in this case the organs from [52]) is:

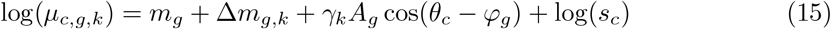

The gene acrophases *φ*_*g*_, amplitudes *A*_*g*_, and basal expression levels *m*_*g*_ are shared across all contexts, while context-specific offsets Δ*m*_*gk*_ and amplitude scaling factors *γ*_*k*_ absorb organ-dependent effects. The shared parameters {*φ*_*g*_, *A*_*g*_, *m*_*g*_} define the *Clock-Reference* (Fig. 1B, Fig. S1A). The fixed gene phases ensure that the meaning of the phases remain the same across context.

#### Cell-type-specific extended gene sets (*Extended-Set*)

To identify rhythmic transcripts for cell-type-specific gene sets, raw expression counts were fit against external collection times using a Generalized Linear Model (GLM) with a single harmonic. Genes were retained for the clock set *G*_*extended*_ according to:

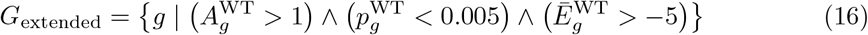

where *A*_*g*_ is the log_2_ fold-change amplitude, *p*_*g*_ is the rhythmicity p-value (Benjamini-Hochberg corrected Likelihood Ratio Test against a flat-line null model), and 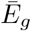 is the log_10_ mean expression.

#### Knockout-based circadian gene filtering

As an alternative strategy independent of collection time, we identified high-confidence clock-driven genes by comparing rhythmic parameters between wild-type (WT) and clock-disrupted (*Bmal1* and *Cry1/Cry2* knockout) conditions [44]. Differentially rhythmic genes were identified using the DryR method, which employs the Bayesian Information Criterion (BIC) to classify genes as rhythmic in WT but arrhythmic in the knockout. We reimplemented DryR in Python as part of the *scRitmo* suite. Candidate genes were further filtered using the criteria in Eq. (16).

#### Statistical analysis

To compare absolute phase errors (AE) across different computational methods, sequencing depths, or hyperparameter settings, we employed the two-tailed Mann-Whitney U test (also known as the Wilcoxon rank-sum test). This non-parametric approach was selected because AE distributions typically exhibit significant positive skewness and include high-error outliers, violating the normality assumptions required for parametric t-tests.

#### Circadian clock comparison across cell types

In (Fig. 3I Fig. 4B) we compare circadian rhythmicity across cell types. To do so we compared the parameters (Eq. (2)) for each cell-type *k* and each gene *g*. To place all cell types on a common reference, we estimated a per-cell-type phase offset Δ_*k*_ by minimising the circular distance between the fitted phases and a reference time vector *t* (ZT or CT). The offset was computed as

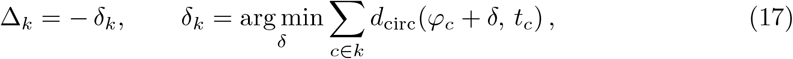

where the sum runs over all cells belonging to cell-type *k* and *d*_circ_ is the circular distance. To quantify the overall rhythmic strength of each cell-type relative to the gene population, we computed an expression-weighted mean of centred amplitudes. For each gene *g* and cell type *k*, the amplitude *A*_*g,k*_ was centred by subtracting the cross-cell-type median *Ã*_*g*_, and the result was weighted by the linear-scale mean expression 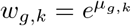:

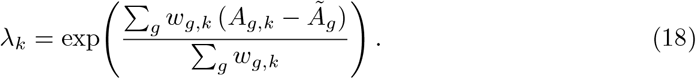

Values *λ*_*k*_ *>* 1 indicate that a cell type exhibits, on average, stronger-than-median circadian amplitudes across the measured gene set, while *λ*_*k*_ *<* 1 indicates weaker rhythmicity. The two quantities Δ_*k*_ and *λ*_*k*_ are jointly visualized in a polar plot, where the angular coordinate encodes the phase offset and the radial coordinate encodes the rhythmicity score.

#### Absolute error and Median absolute error

To quantify the phase inference in simulations, we used used the absolute difference between simulated ground truth phases and inferred phases (Fig. 1 Fig. S1). To evaluate the phase inference in real samples, we used the external collection time as a reference label. While this does not constitute a true ground truth internal phase, cells within a population may exhibit systematic shifts or desynchrony relative to the external time, it remains a widely adopted benchmark in the field and provides a meaningful proxy: in many cases where we expect good synchrony (e.g. in tissues), better alignment between inferred phases and the reference time is the best available indicator of precise inference. To cope with a global phase shifts, for each cell type, the inferred phases were shifted by a global offset, optimised over 200 discretized shifts over the full circle, to minimise the median absolute error (MAE) against the reference phases. This alignment removes any arbitrary global phase offset and ensures the MAE reflects only the spread of individual cells around the reference, not a systematic shift of the population.

*Note:* that in this paper, we have chosen to report all internal phases in units of time, i.e. using CT times between 0 and 24 h rather than less intuitive angles.

### 4.4 Period variance estimation from SABER-FISH data progressive desynchronization

To recover the distribution of single-cell free-running periods from static population snapshots, we modelled the phase evolution of each oscillator as a linear function of time, *θ*(*t*) = *ωt*, where *ω* is the angular velocity. Assuming the population of angular velocities follows a Normal distribution 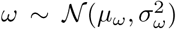, the phase at time *t* is also normally distributed according to:

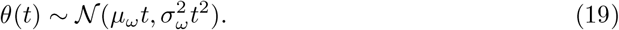

The order parameter *R*(*t*) = 𝔼 [*e*^*iθ*(*t*)^], representing the coherence of the population, is defined as the magnitude of the first circular moment of the phase distribution. The population mean term vanishes upon taking the modulus, yielding a decay profile dependent solely on the variance of the angular velocities:

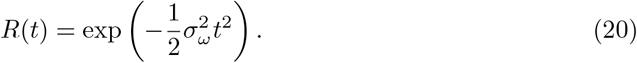

The corresponding circular standard deviation grows linearly:

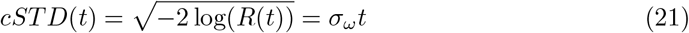

This analytical form allows direct estimation of *σ*_*ω*_ from the slope of the observed cSTD over time. Finally, to interpret this variability in the temporal domain, we derive the distribution of periods *T* = 2*π/ω*. Using the change of variables formula 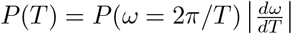 with the Jacobian 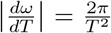, we obtain the probability density function for the periods, which exhibits the characteristic skewness arising from the reciprocal transformation of a Gaussian variable.

### 4.5 Pre-processing of murine scRNA-seq data

#### Alignment

All scRNA-seq datasets were aligned using a Snakemake pipeline built around STARsolo [7], which performs simultaneous alignment, cell barcode demultiplexing, and UMI counting. The pipeline quantifies both exonic and intronic reads, producing spliced and unspliced count matrices. The pipeline is available at https://github.com/cgob/SingleCell_IE_snakemake.

#### Cell filtering

To ensure high-quality transcriptomic profiles for downstream analysis, we performed cell-level filtering based on several quality control metrics. We removed cells with excessively low library sizes to exclude empty droplets or low-quality captures, as well as cells with abnormally high total counts to mitigate the impact of potential doublets. Additionally, cells exhibiting a high proportion of mitochondrial transcripts were excluded, as this typically serves as an indicator of cellular stress or compromised membrane integrity. Finally, we filtered cells based on their unspliced transcript fraction to ensure that we only retained viable, transcriptionally active cells and excluded profiles dominated by ambient mature mRNA or degraded cellular material.

#### Cell-type annotation

Cell type annotation was performed using the scVI (single-cell Variational Inference) framework [23]. Raw spliced counts were used as input, and the model was trained with batch correction across samples (*n*_latent_ = 10, *n*_hidden_ = 128, *n*_layers_ = 1, max epochs=391). The resulting 10-dimensional latent representation was used to compute a nearest-neighbor graph (*k* = 30), followed by Leiden clustering [42] (resolution=0.3). Clusters were manually annotated based on the expression of established marker genes.

### 4.6 Preprocessing of *Drosophila* Clock Neuron scRNA-seq Alignment

Raw sequencing data from the CEL-Seq2 protocol [24] were aligned using STARsolo [7], producing per-cell gene expression count matrices. Exonic and intronic counts were derived separately, with unspliced counts computed as the difference between total (GeneFull) and exonic (Gene) matrices.

#### Cell-type annotation

All preprocessing and clustering were performed using Scanpy [48]. After initial filtering (minimum 100 counts per cell, minimum 100 cells per gene), quality control metrics were computed including mitochondrial content and unspliced transcript fraction. Cells were filtered based on unspliced fraction, mitochondrial content, and total counts. Ribosomal, mitochondrial, and rRNA genes were removed prior to downstream analysis. All QC metrics used are described in Table 1. The 3,000 most variable genes were selected using the Seurat v3 method [38], and batch effects across samples were corrected with scVI [23] (*n*_layers_ = 2, *n*_latent_ = 30, gene likelihood = negative binomial), using per-sample identity as batch key and mitochondrial, ribosomal, and unspliced fractions as continuous covariates. The resulting latent representation was used to build a *k*-nearest-neighbor graph for UMAP embedding and Leiden clustering [42] (resolution = 0.5). Clock neuron populations were identified by filtering Leiden clusters based on pseudobulk expression of canonical clock genes (*tim, vri*), followed by sub-clustering at higher resolution (Leiden, resolution = 1.0). Cell types were annotated using marker genes from the original study [24] and connectomic data [33], including neuropeptides and their receptors (*Pdf, AstC, CNMa, NPF, Trissin, ITP, sNPF, Rh7, VGlut*, among others), yielding 16 clock neuron clusters.

**Table 1:** Cell Filtering Criteria for Single-Cell Datasets.

| Dataset | GEO | Counts/Cell | % Unspliced | % Mitochondrial | Min Cells/Gene |
| --- | --- | --- | --- | --- | --- |
| LIVER [8] | GSE145197 | 1,200–6,000 | 2.5% – 25% | Not filtered | 3 |
| SKIN [10] | GSE223109 | 8,000–50,000 | > 6% | < 13% | 3 |
| AORTA [2] | GSE206583 | 3,000–50,000 | 5% – 35% | < 13% | 3 |
| DROSOPHILA [24] | GSE157504 | 2,000–30,000 | 7.5% – 25% | 1.5% – 15% | 100 |

### 4.7 FLASH-Seq experiment

The NIH3T3-Rev-VNP1-H2B-iRFP dual-reporter cell line was derived from NIH3T3-Rev-VNP1 cells (NIH3T3-Nr1d1-Venus) [27, 3] by lentiviral transduction with the pLenti-H2BiRFP720 construct (Addgene plasmid #128961). Lentiviral particles were produced by co-transfecting HEK293T cells with pLenti-H2B-iRFP720, pMD2.G, and psPAX2. Transduced cells were sorted by FACS to select the iRFP-positive population. Cells were cultured in DMEM high glucose with sodium pyruvate (Gibco) supplemented with 10% fetal bovine serum (FBS) and 1% penicillin-streptomycin. Upon reaching 95% confluency (5 days postseeding), cells were harvested via trypsinization, neutralized with serum-containing media, and centrifuged for 5 minutes. The pellet was washed with PBS and resuspended in FACS buffer (PBS supplemented with 0.5% BSA and 2–5 mM EDTA) at a density of 1 *×* 10^6^ cells/mL. To ensure a single-cell suspension, the sample was filtered through a 40 *µ*m mesh into FACS tubes. DAPI was added immediately prior to sorting to exclude dead cells. Single-cell sorting was performed on a FACS S8 Discovery system. Gating strategies were established to identify the double-positive population using Venus and iRFP channels. To capture the full spectrum of reporter expression, cells were sorted into 384-well plates using a binned strategy: four bins were defined along the YFP (Nr1d1-Venus) axis and two bins along the iRFP (H2B) axis. An equal number of cells was collected for each respective bin to ensure balanced representation across expression levels. Full-length scRNA-seq libraries were generated using MERCURIUS™ FLASH-seq kits (Alithea Genomics). Single cells were sorted into two 384-well plates containing lysis buffer. RNA was denatured at 72°C prior to reverse transcription, followed by cDNA amplification via PCR (21 cycles). The cDNA yield for each well was quantified using the Quant-iT™ PicoGreen™ dsDNA Assay (P11496, Thermo Fisher) on a Varioskan ALF (Thermo Fisher). Purified cDNA was diluted to approximately 200 pg/*µ*L and tagmented. Sequencing indexes and adapters were incorporated through a secondary PCR (8 cycles). The resulting libraries were sequenced on an AVITI system (Element Biosciences, CloudBreak) using paired-end 75 bp reads.

Raw paired-end reads were aligned on a per-cell basis to a custom mouse reference genome (GRCm39, GENCODE vM38) augmented with Venus and iRFP reporter sequences using STAR v2.7.11a [7]. BAM files were sorted by coordinate and indexed with SAMtools. To obtain separate exonic and intronic count matrices, we constructed a combined annotation by extracting exonic intervals from the GENCODE GTF and deriving intronic intervals by subtracting exonic regions from gene body coordinates using BEDTools `subtractBed` [32]. Per-cell exon and intron counts were then quantified from the aligned BAM files using `htseq-count` [1] with `-t exon -m intersection-strict` for exonic reads and `-t intron -m union` for intronic reads, both in unstranded mode. Genes with fewer than 10 total reads across all cells were discarded. Quality-control filtering removed cells based on three criteria: mitochondrial content exceeding 14%, unspliced ratio outside the range [0.11, 0.155], and total spliced counts outside [5 *×* 10^5^, 1.4 *×* 10^6^].

### 4.8 SABER-FISH data

We applied *scRitmo* to the SABER-FISH time-series dataset from [28], specifically using the largest of their datasets, comprising four clock genes (*Bmal1, Nr1d1, Nr1d2, Tef*) measured in NIH3T3 fibroblasts. Consistent with the original study, the earliest time point (6 h post-dexamethasone) was excluded from our analysis, as it likely reflects residual acute transcriptional responses rather than genuine circadian dynamics. Since SABER-FISH quantifies absolute transcript counts for only these four genes, conventional library-size normalization is not applicable. Instead, we used cell area as a proxy for cell size, entering it as the size factor *s*_*c*_ in Eq. (2).

## 5 Discussion

A central principle in circadian biology is that apparent damping of gene expression rhythms at the tissue or population level often does not reflect a loss of oscillatory amplitude within individual cells, but rather the progressive desynchronization of intact, self-sustained oscillators with heterogeneous intrinsic periods or phase dynamics [46, 27, 17]. Despite this well-established concept, accurately quantifying single-cell circadian phase from scRNA-seq data and, crucially, distinguishing biological desynchrony from the technical noise inherent to these measurements has remained an open problem. Previous approaches either lacked rigorous uncertainty quantification (Oscope, reCAT, Cyclum) [18, 22, 21] or provided Bayesian posteriors without explicitly modelling phase heterogeneity at the cell population level (Tempo, VIST) [2, 49]. As a result, the field has been unable to determine how much of the apparent circadian phase spread in a population of single cells reflects genuine intercellular heterogeneity versus measurement artifact.

Here, we introduced *scRitmo*, an unsupervised probabilistic framework that jointly provides a circadian phase estimate and a posterior uncertainty for each cell, and decomposes observed population phase variance into biological and technical components via simulation-calibrated variance subtraction. We validated this decomposition against both synthetic data with known ground-truth desynchrony and SABER-FISH time-series data from synchronized fibroblasts progressively drifting out of phase, providing a principled ground-truth validation of biological desynchrony estimation from transcriptomic data.

### 5.1 Single-cell circadian phase inference with principled uncertainty quantification

*scRitmo* provides robust single-cell circadian phase inference by combining a Negative Binomial count model with a full posterior phase distribution for each cell, yielding both a point estimate and a natural measure of confidence. The posterior uncertainty *σ*_*u*_ consistently predicts inference quality across simulations and diverse biological datasets, enabling principled quality control without external time labels. In unsynchronized 3T3 fibroblasts equipped with a fluorescent circadian reporter, and profiled by deep plate-based scRNA-seq, the inferred phases accurately reconstructed oscillations at both the endogenous mRNA level and the reporter protein level. With the predicted single-cell phases, genome-wide harmonic regression identified over 100 rhythmic genes beyond the input set, demonstrating that *scRitmo* enables transcriptome-wide circadian analysis from a single unsynchronized cell population.

A practical challenge for phase inference in droplet-based scRNA-seq is the low sequencing depth typical of current protocols, which limits the information content of the core clock genes alone. The non-uniform distribution of clock gene peak phases creates a bias towards an attractor phase at low sequencing depths. This can be mitigated by principled expansion of the gene set, either through pseudobulk-derived extended sets or knockout-validated circadian genes [44]. Constructing extended and biologically relevant input gene sets currently requires external time labels or curated knockout data. Unsupervised approaches have been proposed [2], but this introduces a trade-off between statistical stability and mechanistic specificity: inclusion of system-driven, in addition to clock-controlled rhythmic transcripts, may enhance the ability to infer precise phases, with the potential caveat of partially decoupling the inferred phase from genuine circadian clock dynamics. The use of knockout-validated gene sets partially mitigates this concern, but does not eliminate it, as the number and phase diversity of *bona fide* clock-controlled transcripts may be limiting in some tissues.

Importantly, as the sensitivity increases and cost of scRNA-seq continues to decline, higher per-cell sequencing depth is becoming routine. Our simulation and experimental results show that at sufficient depth (*>*20,000 UMI/cell), the *Clock-Reference* alone provides accurate and unbiased phase inference without the need for extended gene sets. This suggests the current gene set constraint is a practical limitation rather than a fundamental one.

### 5.2 Quantifying biological desynchrony from transcriptomic data

By decomposing observed phase variance into biological and technical components through simulation-calibrated variance subtraction, *scRitmo* provides a new transcriptomics-based approach for estimating intercellular circadian phase coherence. Until now, quantitative desynchrony measurements at the single-cell level have relied on real-time bioluminescence or fluorescence reporters [46, 27, 17], which are typically performed in engineered cell lines, while it is harder to capture desynchrony of cells in intact tissues [36]. We validated the decomposition through two independent lines of evidence: in SABER-FISH time-series data, *σ*_bio_ increased monotonically after synchronization while *σ*_technical_ remained constant; in *Drosophila* clock neurons, *σ*_bio_ increased under constant darkness relative to light-dark entrainment.

However, the accuracy of the decomposition depends on how faithfully the generative model captures the various sources of gene expression and RNA count variability. In our model, the sources of heterogeneity that are not directly related to the circadian clock, such as heterogeneity in metabolic state, spatial zonation, or cell cycle phase, can be absorbed into either the parameters of the NB noise model or gene rhythmicity coefficients, hence biasing *σ*_bio_ estimates. For example, particularly at low sequencing depth or with an insufficiently informative gene set, the estimated *σ*_technical_ can exceed *σ*_data_, rendering the subtraction undefined. Future extensions incorporating waveform flexibility and explicit modelling of additional latent variables could refine these estimates. While absolute estimates of *σ*_bio_ can be challenging, the most robust application of the framework consists of *relative* comparisons of *σ*_bio_ across conditions within the same cell type and experimental setup. In this mode, systematic biases in the variance decomposition will typically cancel, and changes in desynchrony can be attributed with confidence to the experimental perturbation. Both validations presented here, the temporal progression post-dexamethasone and the LD-to-DD comparison per neuronal cluster, exemplify this comparative design.

### 5.3 Light-dependent reorganization of the *Drosophila* clock neuron network

Application of *scRitmo* to *Drosophila* clock neurons under Light-Dark (LD) and Constant Darkness (DD) conditions reveals condition-dependent and cluster-specific phase reorganization without widespread loss of oscillation amplitude. Pacemaker neurons (s-LNv and l-LNv) maintained stable phase distributions under DD, consistent with their established role in sustaining free-running rhythms [37, 12]. In contrast, DN1p subclusters exhibited pronounced phase advances and increased desynchrony, suggesting stronger dependence on photic input and network-mediated coupling. The magnitude and cluster-specificity of this LD-to-DD reorganization highlight differential sensitivity to entrainment across neuronal subpopulations and align with emerging connectomic and transcriptomic evidence of functional specialization within the clock network [35, 33, 25]. Rather than damping rhythmic output, removal of light cues appears to reorganize network phase relationships while preserving cell-autonomous oscillatory amplitude. In the analysed dataset, cells were pooled from multiple animals at each time point. The increased *σ*_bio_ under DD across all cell types also likely reflects inter-individual phase drift, for example as a results of animal specific free-running periods, in addition to desynchrony among neurons within a single brain. Disentangling intra-from inter-individual contributions to phase dispersion will require future experiments with single-animal resolution.

### 5.4 Conclusion

*scRitmo* transforms a series of snapshot-like scRNA-seq measurements into a quantitative framework for measuring circadian phase coherence at single-cell resolution. By combining probabilistic phase inference with principled uncertainty quantification and simulation-calibrated variance decomposition, it enables direct separation of biological desynchrony from technical noise. Across mammalian tissues, cultured fibroblasts, and *Drosophila* clock neurons, our results support that population-level amplitude damping predominantly reflects desynchronization of intact oscillators rather than loss of cell-autonomous rhythmicity. Making this distinction quantitative, particularly through relative comparisons across conditions, provides a foundation for probing how coupling, entrainment, and disease reshape circadian organization at the cellular scale. As sequencing depth increases, for example as plate-based experimental designs become more accessible, we anticipate that systematic profiling of intra-tissue desynchrony across pathological contexts such as metabolic disease, chronic inflammation, or ageing [4] will reveal how chronodisruption manifests at the cellular level.

## Author Contributions

Conception or design of the work: A.S., C.G., Y.P., and F.N. Acquisition, analysis, or interpretation of data for the work: A.S., C.G., and Y.P. V.H. designed and performed the FLASH-seq experiment including FACS sorting and library generation. Drafting the work or revising it critically for important intellectual content: A.S. and F.N.

## Data availability

The FLASH-seq data are available on Gene Expression Omnibus (GEO) under accession number GSE325045.

## Code availability

The scRitmo package including phase inference, phase variance decomposition, fit of cycler genes, python version of dryR, and much more functionalities are in https://github.com/AndreaSalati/scRITMO

## Use of Large Language Models

Portions of the code used in this study were generated using large language models and subsequently reviewed and modified by the authors. Additionally, these tools were employed to enhance the clarity, coherence, and linguistic quality of the manuscript. Following this process, the authors reviewed and edited the content to ensure accuracy and take full responsibility for the final publication.

## 5.5 Acknowledgments

We thank Eliane Duperrex for generating the NIH3T3-Rev-VNP1-H2B-iRFP dual-reporter cell line and Maxine Leonardi for producing the lentiviral particles used for transduction. We thank Kevin Blackney from the Flow Cytometry Facility (UNIL, AGORA) for the single-cell sorting. The work was supported by the Swiss National Science Foundation project grant 310030B 201267 to F.N.

## 6 Supplementary Figures

**Figure S1:**
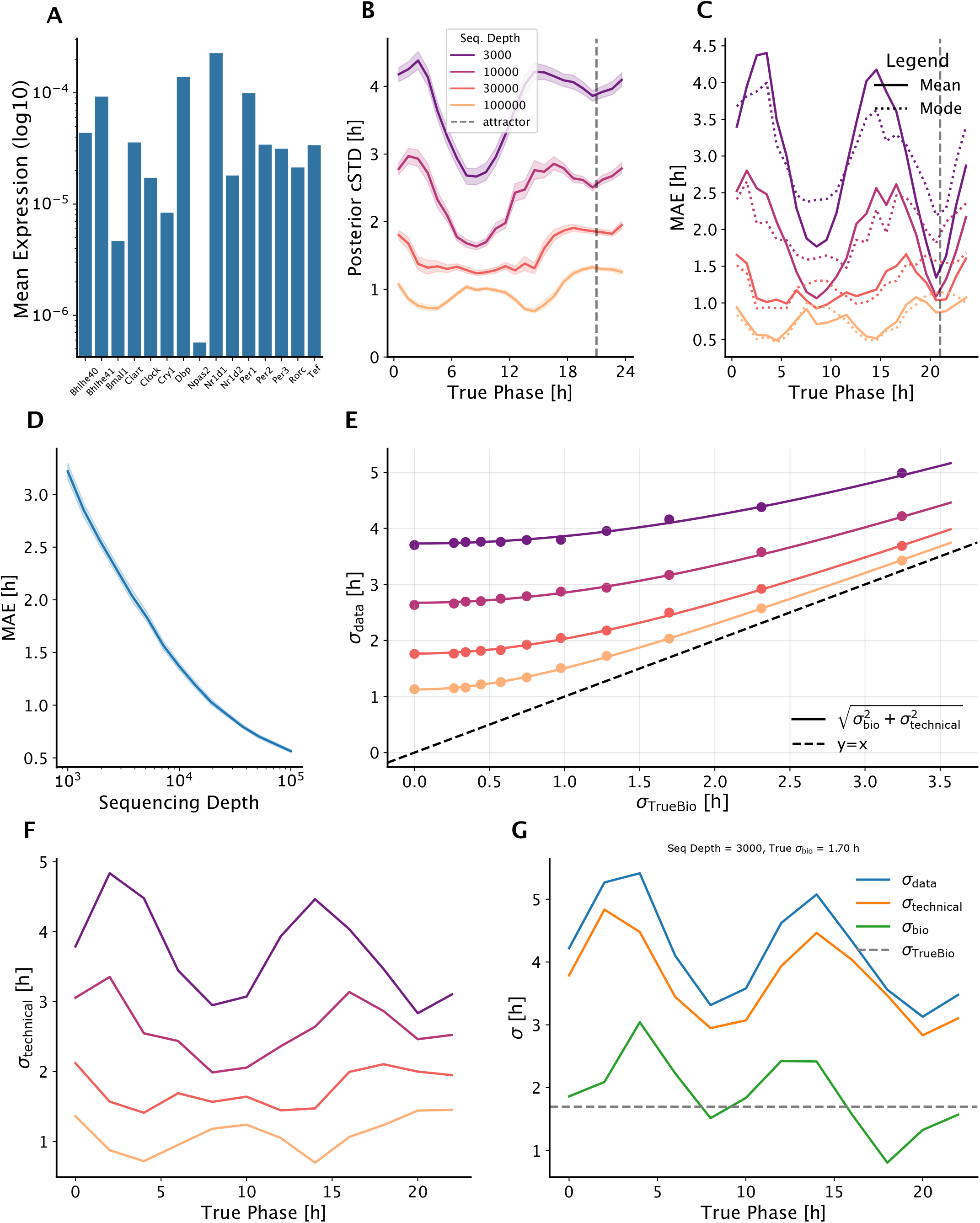
Validation of variance decomposition and phase estimation performance using synthetic data. **A**. Expression baselines of the *Clock-Reference* set, representing transcript counts normalized to a library size of 1. *Clock-Reference* was estimated by fitting a consensus harmonic model across organs from bulk time-series RNA-seq data [52]. **B**. Single-cell posterior circular standard deviation (cSTD) as a function of true circadian phase across varying sequencing depths (median total UMI/cell). **C**. Comparison of phase point-estimation strategies: Median Absolute Error (MAE) using the circular mean (solid lines) versus the posterior mode (dotted lines) across the 24h cycle at varying sequencing depths (median total UMI/cell). **D**. Median absolute error (MAE) of phase estimation as a function of sequencing depth. The solid line indicates the median value while the shaded area represents the 95% confidence interval. **E**. Observed circular standard deviation (*σ*_data_) as a function of ground truth biological desynchrony (*σ*_TrueBio_). The curves illustrate the theoretical relationship 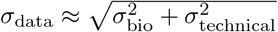, where the y-intercept represents the baseline technical noise (*σ*_technical_). **F**. Estimated technical variability (*σ*_technical_) as a function of true circadian phase across sequencing depths. **G**. Temporal profiles of variance components (*σ*_data_, *σ*_technical_, and inferred *σ*_bio_) compared to the ground-truth biological desynchrony *σ*_TrueBio_ (dashed grey line). Sequencing depth = 3’000 (median total UMI/cell), *σ*_TrueBio_ = 1.7 hours.

**Figure S2:**
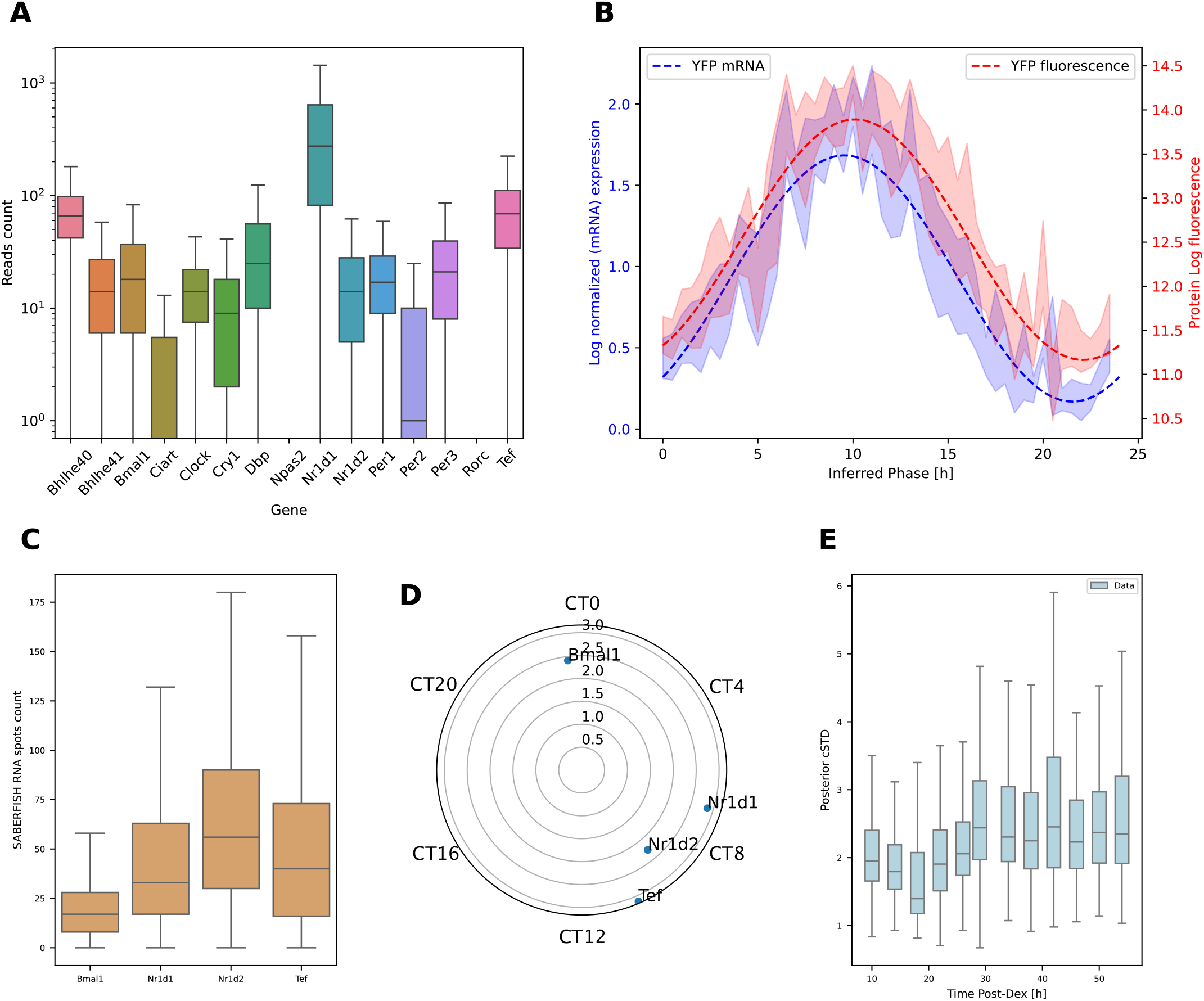
Validation of *scRitmo* phase inference using protein levels and SABER-FISH data. **A–B**. Plate-based scRNA-seq (FLASH-seq) data in unsynchronized 3T3 fibroblasts with simultaneous FACS-based quantification of a Nr1d1-Venus (Rev-Erb*α*-YFP) circadian reporter. **A**. Boxplot showing the distribution of RNA read counts for individual core circadian clock genes in the FLASH-seq dataset. **B**. Phase-resolved comparison between *YFP* mRNA expression (blue) and corresponding YFP protein fluorescence intensity (red) in FLASH-seq data. Dotted lines represent harmonic fits to the data, while shaded areas indicate the 95% confidence interval. **C–E**. SABER-FISH time-series data post-dexamethasone (Dex) shock in 3T3 fibroblasts [28]. **C**. Boxplot of the absolute number of mRNA molecules per cell for *Bmal1, Nr1d1, Nr1d2*, and *Tef* as measured by SABER-FISH. **D**. Polar plot displaying the rhythmic parameters for the four genes in (C) as estimated by *scRitmo*. The phases are rotated to align *Bmal1* at CT23. **E**. Boxplot of the posterior circular standard deviation (cSTD) across individual cells as a function of time post-dexamethasone treatment in the SABER-FISH dataset.

**Figure S3:**
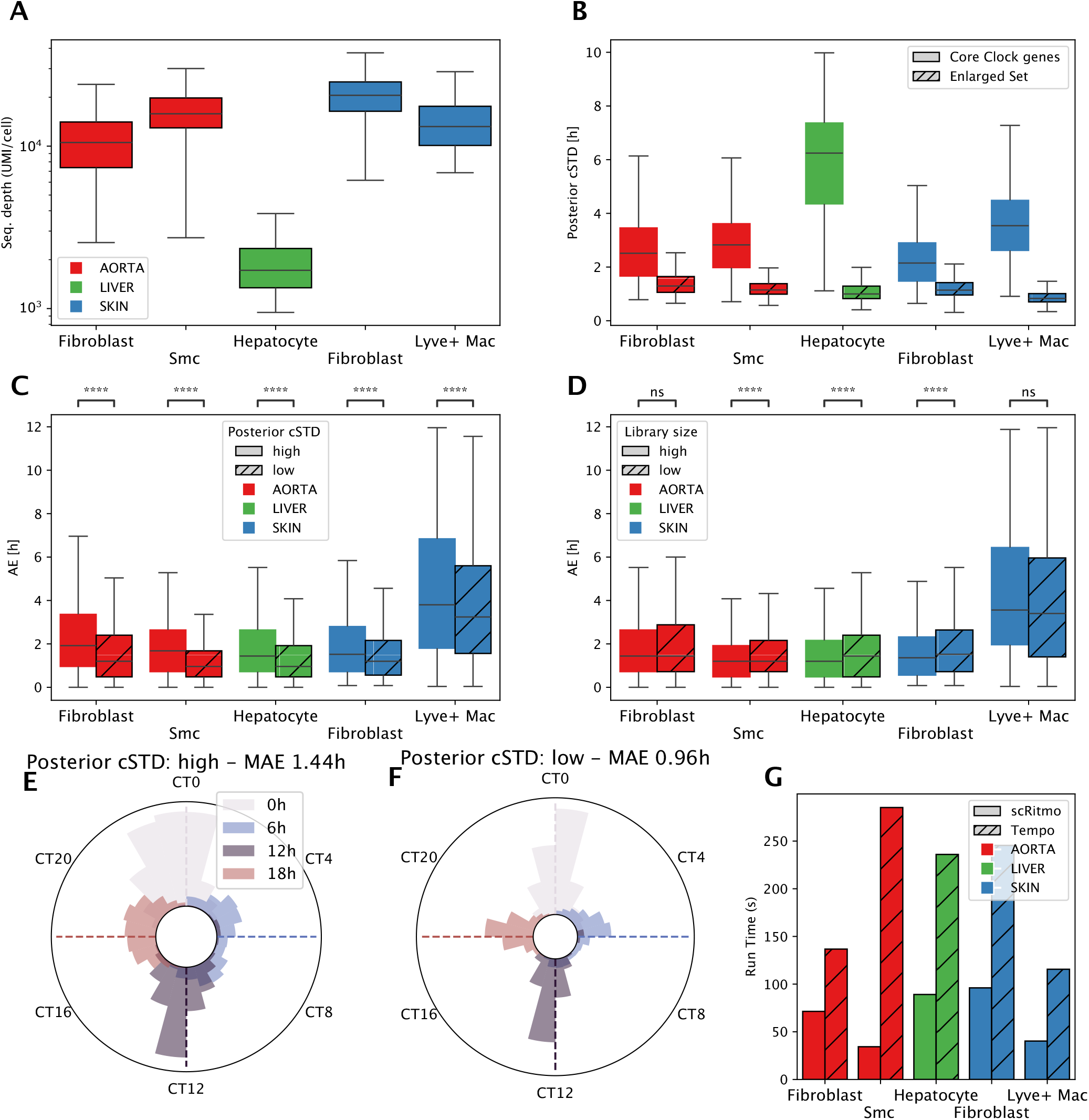
Technical validation and benchmarking of single-cell circadian phase inference across mammalian tissues. **A**. Box plots displaying the sequencing depth (UMI per cell) for the analyzed cell types across aorta, liver, and skin tissues. **B**. Boxplots of posterior circular standard deviations. Solid is *Clock-Reference* while hatched is *Extended-Set*. **C**. Box plots of phase estimation errors stratified by posterior uncertainty (higher versus lower than median posterior *σ*_*u*_) across cell types. Statistical significance for all pairs was determined by two-tailed Mann-Whitney U tests (****, *P <* 0.0001). **D**. Box plots of phase estimation errors stratified by library size (high versus low) across cell types. Statistical testing used is the same as in (D). **E**,**F**. Polar histograms illustrating the distribution of inferred phases for cells exhibiting high (E) and low (F) posterior uncertainty in liver hepatocytes. *Extended-Set* was used for phase inference. Histograms are colored by ground truth timepoints (0h, 6h, 12h, 18h). Dashed lines indicate the circular ZT collection time. **G**. Bar chart comparing the computational run time in seconds between *scRitmo* and Tempo across different cell types. Solid is *scRitmo* while hatched is Tempo.

**Figure S4:**
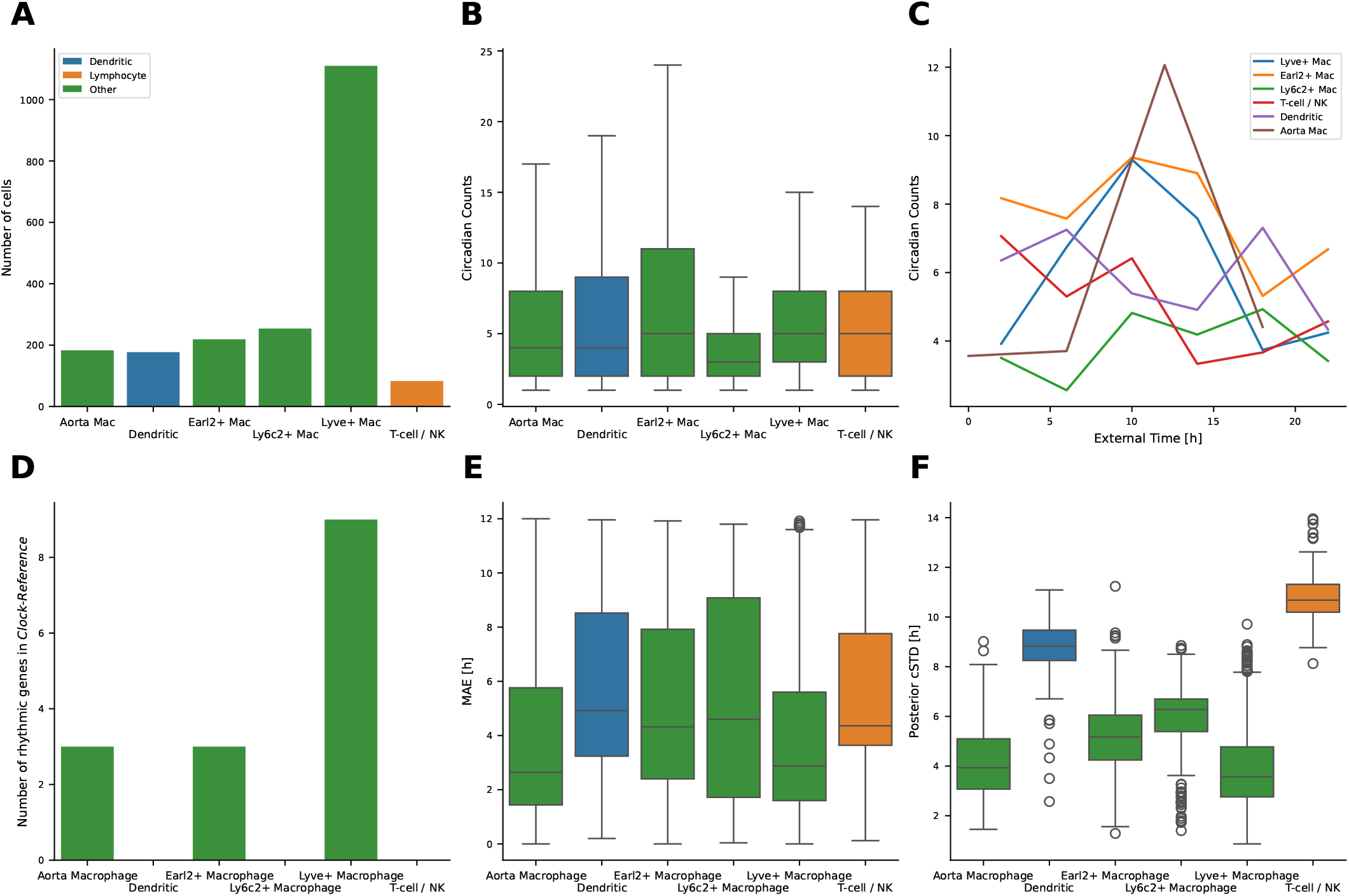
Heterogeneity and inference uncertainty of circadian rhythms across aortic and skin immune cells. **A**. Total cell counts for each immune cell type analyzed from aorta and skin datasets. **B**. Distribution of circadian counts (sum of 15 core clock gene counts) across cell types. **C**. Temporal profiles of circadian counts across external time points. Only Lyve^+^ macrophages and aortic macrophages display the expected canonical distribution peaked around 10 h (corresponding to the *Dbp* expression peak), while other cell types show disrupted or arrhythmic patterns. **D**. Number of significantly rhythmic core clock genes per cell type after fitting expression to external time (filtered by amplitude, statistical significance, and mean expression). **E**. Median absolute error (MAE) between inferred single-cell circadian phases and external reference time, (*Clock-Reference* was used). Only two cell types exhibit MAE *<* 3 h. **F**. Posterior standard deviations (uncertainty) of single-cell phase estimates. Non-macrophage cell types (dendritic cells and T-cells/NK cells) display substantially higher inference uncertainties compared to macrophage populations, consistent with weaker or incoherent clock gene expression.

**Figure S5:**
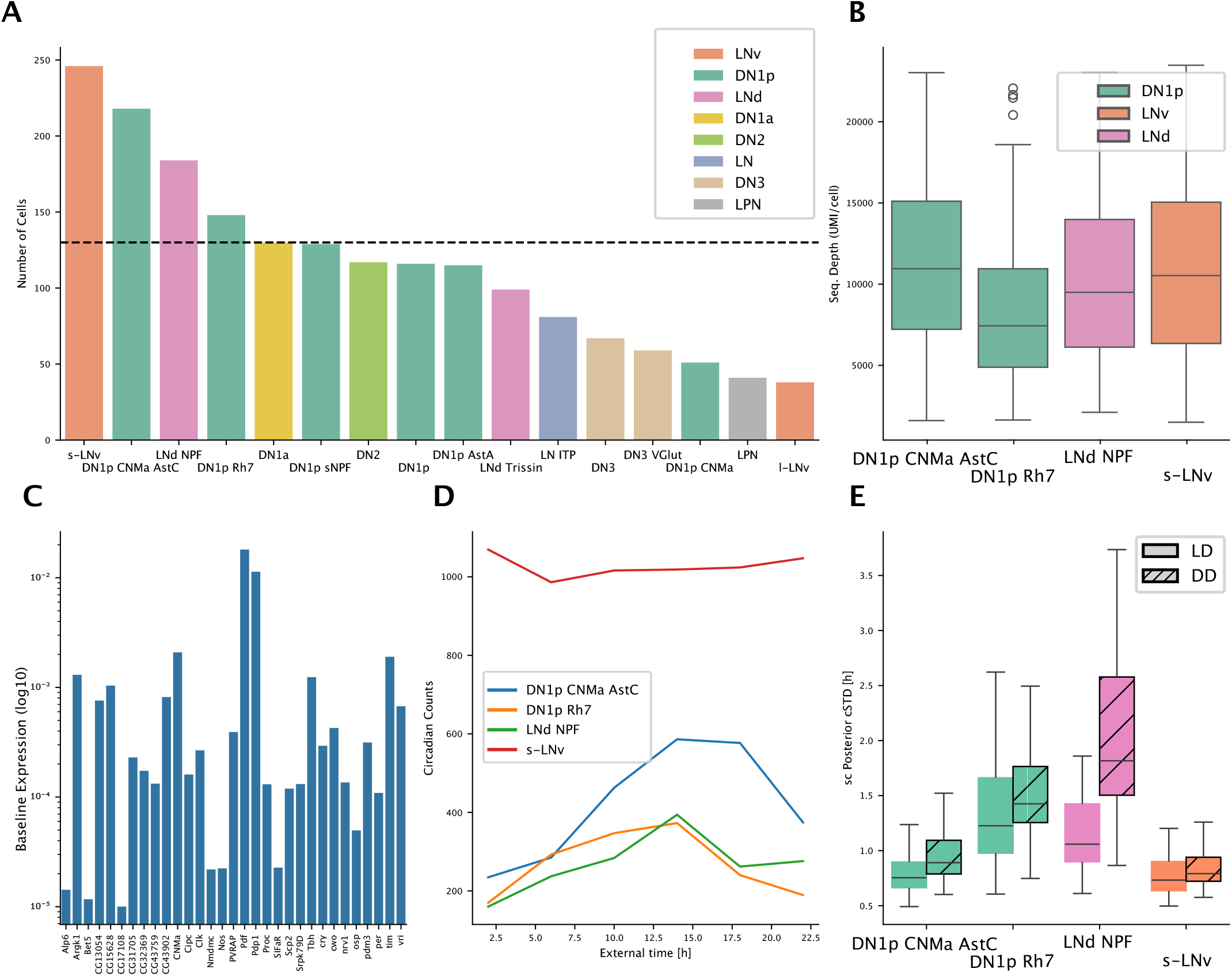
Quality control and parameter estimation for Drosophila clock neurons. **A**. Bar plot showing the number of cells per cell type cluster. A threshold of 130 cells (dashed line) was applied to ensure sufficient data across experimental conditions and timepoints for subsequent analysis. **B**. Boxplots of sequencing depth (UMI counts per cell) for the major cell type clusters. **C**. Baseline expression parameters (log10 scale) for the *Drosophila-Set* genes as estimated by the *scRitmo* model **D**. Aggregate circadian transcript counts over external time, showing the cumulative expression profiles of the *Drosophila-Set* across the four major clock neuron clusters. **E**. Boxplots of the single-cell posterior circular standard deviation (cSTD) for major cell types under entrained (LD, solid) and free-running (DD, hatched) conditions.

